# Genomic Dissection of an Icelandic Epidemic of Equine Respiratory Disease

**DOI:** 10.1101/059949

**Authors:** Sigríður Björnsdóttir, Simon R. Harris, Vilhjálmur Svansson, Eggert Gunnarsson, Ólöf G. Sigurðardóttirr, Kristina Gammeljord, Karen F. Steward, J. Richard Newton, Carl Robinson, Amelia R. L. Charbonneau, Julian Parkhill, Matthew T.G. Holden, Andrew S. Waller

**Author notes:** Correspondence should be addressed to A.S.W (; [).].

## Abstract

The native horse population of Iceland has remained free of major infectious diseases. Between May and July 2010 an epidemic of respiratory disease swept through the population. Initial microbiological investigations ruled out known equine viral agents as the cause of the infections, but identified the opportunistic pathogen *Streptococcus zooepidemicus* as being frequently isolated from diseased animals. This diverse bacterial species has a broad host range and is usually regarded as a commensal of horses. By genome sequencing *S. zooepidemicus* recovered from horses during the epidemic we show that although multiple clones of *S. zooepidemicus* were present in the population, one particular clone, ST209, was responsible for the epidemic. Concurrent with the epidemic, ST209 caused zoonotic infections, highlighting the pathogenic potential of this clone. Phylogenetic analysis suggests that the original ST209 strain entered Iceland in late 2008 or early 2009. Epidemiological investigation revealed that the incursion of this strain into a training yard that utilized a submerged treadmill between the 5^th^ and 19^th^ of February 2010 was a critical trigger for the ensuing epidemic of disease, provided a nidus for the infection of multiple horses, and subsequent distribution of these animals to multiple sites in Iceland.

## INTRODUCTION

The Icelandic horse population is geographically isolated and arose from animals introduced by settlers in the 9^th^ and 10^th^ centuries, with virtually no import of horses for the last 1,000 years. Icelandic horses are extremely hardy and are prized for their maneuverability and movement; Icelandic horses possess five distinct gaits, instead the three gaits common to most other horses. The isolation of the population has meant that Icelandic horses have remained free from most common contagious diseases of Equidae including equine influenza, equine rhinopneumonitis, equine viral arteritis, *Rhodococcus equi* and strangles (*Streptococcus equi* subsp. *equi*) (Robinson, Steward, Potts, Barker, Hammond, Pierce, Gunnarsson, Svansson, Slater, Newton et al. 2013; Sanz, Oliveira, Loynachan, Page, Svansson, Giguere and Horohov 2014; Torfason, Thorsteinsdottir, Torsteinsdottir and Svansson 2008). As such the population is extremely vulnerable to these and other infectious agents and strict biosecurity measures are in place to maintain a high health status.

Between April and November of 2010, an epidemic of respiratory disease that was typified by a persistent dry cough and mucopurulent nasal discharge, swept across Iceland affecting almost the entire population of an estimated 77,000 horses. In addition to the paralysis of all equine activities it resulted in a self-imposed ban on the export of horses; with important economic consequences for the Icelandic horse industry.

Here we describe how the opportunistic bacterial pathogen *Streptococcus equi* subsp. *zooepidemicus* (*S. zooepidemicus*) was identified as the causative agent of the respiratory disease and how, by analyzing the genome sequences of *S. zooepidemicus* isolates, a distinct rapidly expanding causal clone was identified. Furthermore, we demonstrate that this clone was responsible for zoonotic infections during the course of the epidemic, and that the epidemic strain has become endemic within the Icelandic horse population.

## RESULTS

### Epidemiology of a respiratory disease epidemic

The first cases of the epidemic were reported on the 7th of April at an equine center in the north of Iceland (farm 16). At that time, 10 out of the 42 horses showed clinical signs of respiratory tract infection, one of which had been coughing for 16 days. Three other horses at farm 16 had a history of similar clinical signs two to six weeks earlier. Information received from other parts of the country within that week, revealed a widespread infection throughout the country and it became apparent that an epidemic could not be avoided.

Dry coughing was usually the first clinical sign noted; most often coexisting with mucopurulent nasal discharge and mild conjunctivitis, although rectal temperature remained normal in most horses (Supplementary Table 1). Laryngoscopy revealed laryngitis and inflammation of the upper trachea. The incubation period was between two and four weeks and the duration of clinical signs varied from two to ten weeks. The entire equine population in Iceland appeared to be susceptible to the disease resulting in almost 100% morbidity, but very low mortality.

### Identification of *S. zooepidemicus* as the causative agent

The initial assumption due to the rate of spread was that a virus was responsible. Nasal swabs were tested by PCR for viruses known to cause respiratory disease in horses and some common respiratory viruses of humans and animals (Supplementary Table 2). Paired blood samples were used to determine if horses developed an antibody response to equine respiratory viruses during the course of their disease. Established and primary cell lines were also used in attempts to isolate a novel causal virus. All of these tests proved negative with the exception of equine gammaherpesviruses, which were isolated from small numbers of both healthy and clinically affected horses and therefore these viruses were not regarded as being related to the epidemic.

In the absence of a viral pathogen, it was noted that the Gram-positive bacterium *S. zooepidemicus* was isolated from almost all of the nasal swabs taken from coughing horses and from the diseased tissues of occasional fatal cases (Supplementary Table 1). Initially this was not considered of relevance, as although *S. zooepidemicus* is an opportunistic pathogen associated with a range of equine infections (Wood, Newton, Chanter and Mumford 2005a; Wood, Newton, Chanter and Mumford 2005b) and infections of other animals including humans (Abbott, Acke, Khan, Muldoon, Markey, Pinilla, Leonard 2010, Steward and Waller 2010; Balter, Benin, Pinto, Teixeira, Alvim, Luna, Jackson, LaClaire, Elliott, Facklam et al. 2000), it is routinely isolated from healthy horses and is widely considered to be a commensal. Evidence for the role of *S. zooepidemicus* in the outbreak was provided when three healthy horses (horses 16, 17 and 18, Supplementary Table 1) from the University of Iceland were transferred into a barn (farm 5) on the 14^th^ June. The seventeen resident horses had been diagnosed with respiratory disease on the 31^st^ May and *S. zooepidemicus* was recovered from the nasal swabs of eleven horses, which were sampled on the 2^nd^ June. Following their introduction, the three healthy horses were monitored for clinical signs of disease. Ten days post-arrival, nasal discharge was apparent, which became mucopurulent 9 days later accompanied by the first observation of coughing. *S. zooepidemicus* was recovered from these horses from 20 days post-introduction. Post-mortem examination of these horses revealed signs of respiratory infection, which included the presence of mucopurulent material in the nasal cavity, larynx and trachea and enlarged cervical and mandibular lymph nodes. Histopathological analysis identified sub-acute rhinitis, laryngitis and tracheitis with transmigration of neutrophils through the mucosal epithelium. There was a mild hyperplasia and metaplasia of the respiratory epithelium of the trachea in all three horses, with loss of goblet and ciliated cells and focal erosions. Lympho-histiocytic and plasmacytic inflammation was seen in the superficial submucosa, forming a broad inflammatory band in the trachea of horse 17 and in the nasal mucosa of horse 16. *S. zooepidemicus* was isolated in large numbers from the nasal cavity, larynx and trachea from all three horses, and also isolated from the nasopharynx of horses 16 and 18, from a bronchus of horse 16, and from the guttural pouch of horse 18.

### Genomic investigation of the outbreak

To determine if the epidemic was associated with the introduction and spread of a specific *S. zooepidemicus* strain, we employed whole genome sequencing (WGS) to interrogate the relationships of 251 *S. zooepidemicus* isolates recovered from the nasal swabs of horses that were either actively showing signs of respiratory disease or were stabled on affected premises during the course of the epidemic (Supplementary Table 1). Included in this were multiple isolates recovered from the same clinical sample or the same horse sampled over time, which optimized the isolation of the epidemic strain in the face of concomitant colonization. During the epidemic, cases of *S. zooepidemicus* disease were reported in companion animals (two cats and one dog) and three people and these additional isolates were included in our analysis. In order to provide a wider temporal and genetic context, we included sequenced Icelandic isolates from seven horses, one dog and two sheep that predated the 2010 epidemic, and also included the genomes of 38 strains of *S. zooepidemicus* from outside Iceland that were representative of the known species diversity as a whole as defined by multilocus sequence typing (MLST) (Webb, Jolley, Mitchell, Robinson, Newton, Maiden and Waller 2008) (Supplementary Table 1).

### Population structure of Icelandic *S. zooepidemicus* is dominated by a few dominant clones

The availability of MLST has permitted the identification of strains that were associated with specific infections, including respiratory disease and abortion in horses and acute fatal haemorrhagic pneumonia in dogs, suggesting that certain strains of *S. zooepidemicus* may have greater potential to cause disease (Abbott 2010, Acke, Khan, Muldoon, Markey, Pinilla, Leonard, Steward and Waller 2010; Chalker, Waller, Webb, Spearing, Crosse, Brownlie and Erles 2012; Webb, Jolley, Mitchell, Robinson, Newton, Maiden and Waller 2008). However, the identification of specific disease-causing strains can be confounded by the limited ability of MLST to differentiate closely related strains. Phylogenetic analysis of the WGS data provided a high-resolution view of the Icelandic *S. zooepidemicus* population structure. The majority of *S. zooepidemicus* recovered during the epidemic (201 of 257 isolates (78%) from 33 premises) fell into four distinct clades (Supplementary Table 1 and Figure 1) that corresponded to four separate MLST sequence types (STs): Clade 4 (ST306) contained 37 isolates obtained from the Institute for Experimental Pathology at Keldur, Reykjavik between September and November 2010 and differed by a maximum of 44 SNPs; Clade 3 (ST248) contained 52 isolates obtained from 8 different farms, and a canine isolate, and differed by a maximum of 151 SNPs; Clade 2 (ST246) contained 29 isolates, obtained from 10 farms, and a human isolate; and differed by a maximum of 153 SNPs; and Clade 1 (ST209) containing 83 isolates from 22 farms, and a human and a feline isolate that differed by a maximum of 25 SNPs.

**Figure 1.**
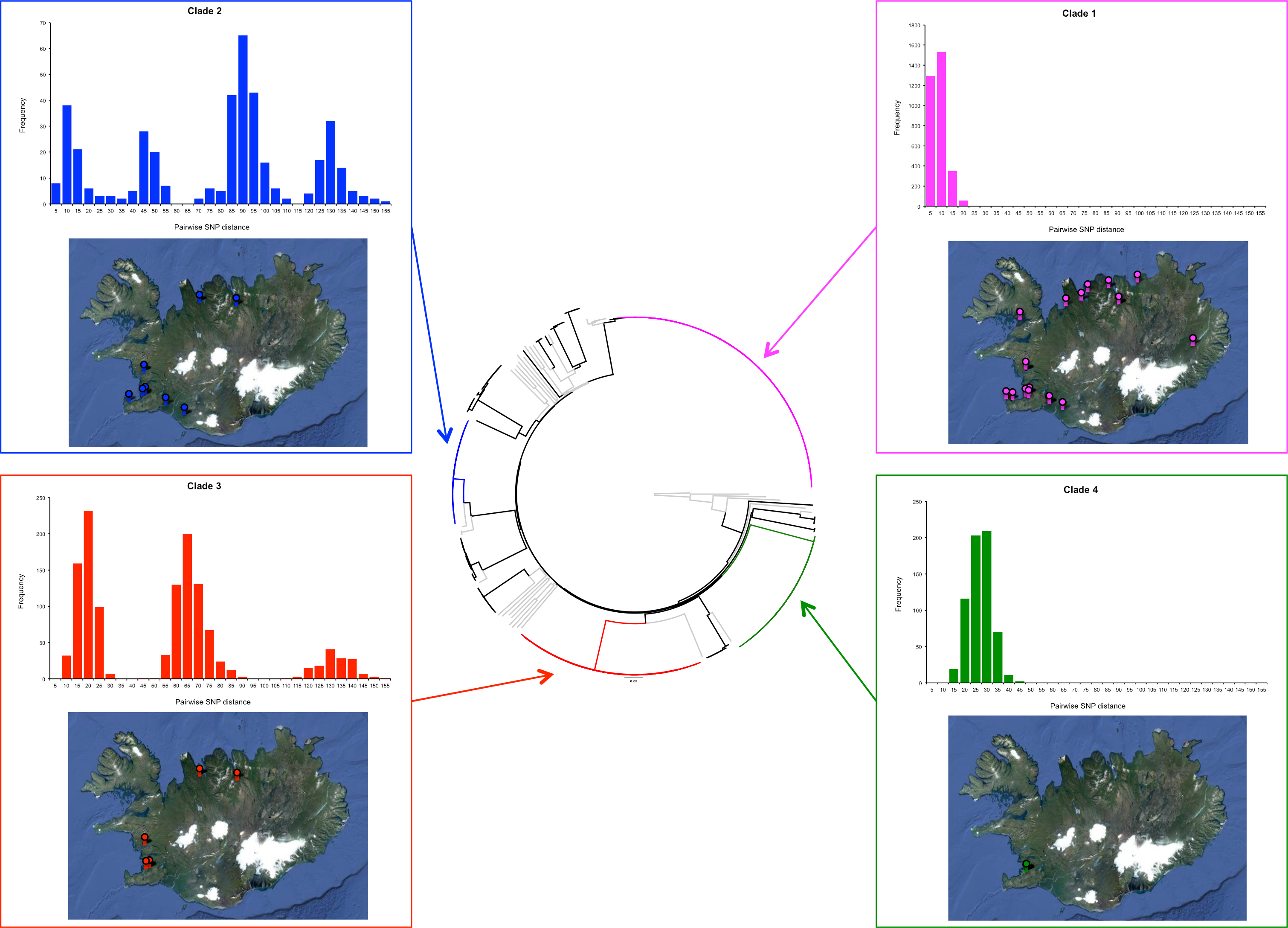
Diversity of the Iceland *S. zooepidemicus* population. Phylogenetic reconstruction of the Icelandic *S. zooepidemicus* population structure (center); Neighbor Joining phylogenetic tree was built using core SNPs, with SNPs in regions of recombination removed. Included in the phylogeny were *S. zooepidemicus* isolates from outside Iceland that were representative of the genetic diversity of the species. The branches of the tree containing these isolates are colored in grey. The four main clades in the Iceland population: clades 1, 2, 3, and 4, are colored magenta, green, red and blue respectively. For each of the clades the distribution of the pairwise SNPs distance of calculated for the core genome are displayed (top graph in panel), and also the geographic distribution of the isolates’ origins within Iceland (bottom image in panel).

### Confirmation that the ST209 (clade 1) *S. zooepidemicus* population was the cause of the epidemic

The wide geographic dispersal of ST209 and its relative lack of diversity indicate that this strain had been transmitted to horses throughout Iceland in a short period of time, suggesting that this strain was responsible for the epidemic of respiratory disease. Evidence to support this came from the investigation of the healthy horses that were introduced to farm 5 and subsequently acquired disease. Seven of the 14 isolates recovered from these horses were ST209, whilst one was ST246. Five of the six remaining isolates that were recovered from these horses clustered together as a separate group within the Icelandic population of *S. zooepidemicus* and were not recovered from horses that were already resident on farm 5 (Supplementary Table 1). One of the horses introduced into farm 5 (horse 17) did not provide a ST209 strain. Notably this horse was euthanized 23 days post-introduction in contrast to the other two horses that were euthanized 28 days post-introduction. A possible explanation for the absence of ST209 may be that these strains were not shed in any great number until the later time point, and the heterogeneous population of *S. zooepidemicus* may have confound the identification of the epidemic strain during the early stages of infection. In this regard it is worth noting that all of the isolates recovered from nasal swabs taken from horses 16 and 18 from day 28 and at the time of post-mortem examination on day 30 belonged to ST209.

Further evidence of the localized transmission of ST209 can be found in the clustered regularly interspaced short palindromic repeats (CRISPRs), which provides a snapshot of a strain’s exposure to mobile genetic elements (Holden, Heather, Paillot, Steward, Webb, Ainslie, Jourdan, Bason, Holroyd, Mungall et al. 2009; Waller and Robinson 2013). In keeping with rapid transmission across the population, the Icelandic isolates of ST209 predominantly shared the same complement of 41 spacer sequences (Figure 2). However, ST209 isolates from horses on farm 5 were found to contain a novel spacer 35 sequence. This unique spacer was present in isolates recovered from four resident horses (3, 5, 7 and 14), but not from three other residents (1, 2 and 9) on the 2^nd^ June, or from any other affected horses from other farms throughout Iceland. The ST209 isolates that were recovered from the horses, which were introduced to farm 5 (16 and 18) contained the unique spacer 35 sequence directly linking the acquisition of this strain to their arrival and exposure to the resident horses at farm 5.

**Figure 2.**
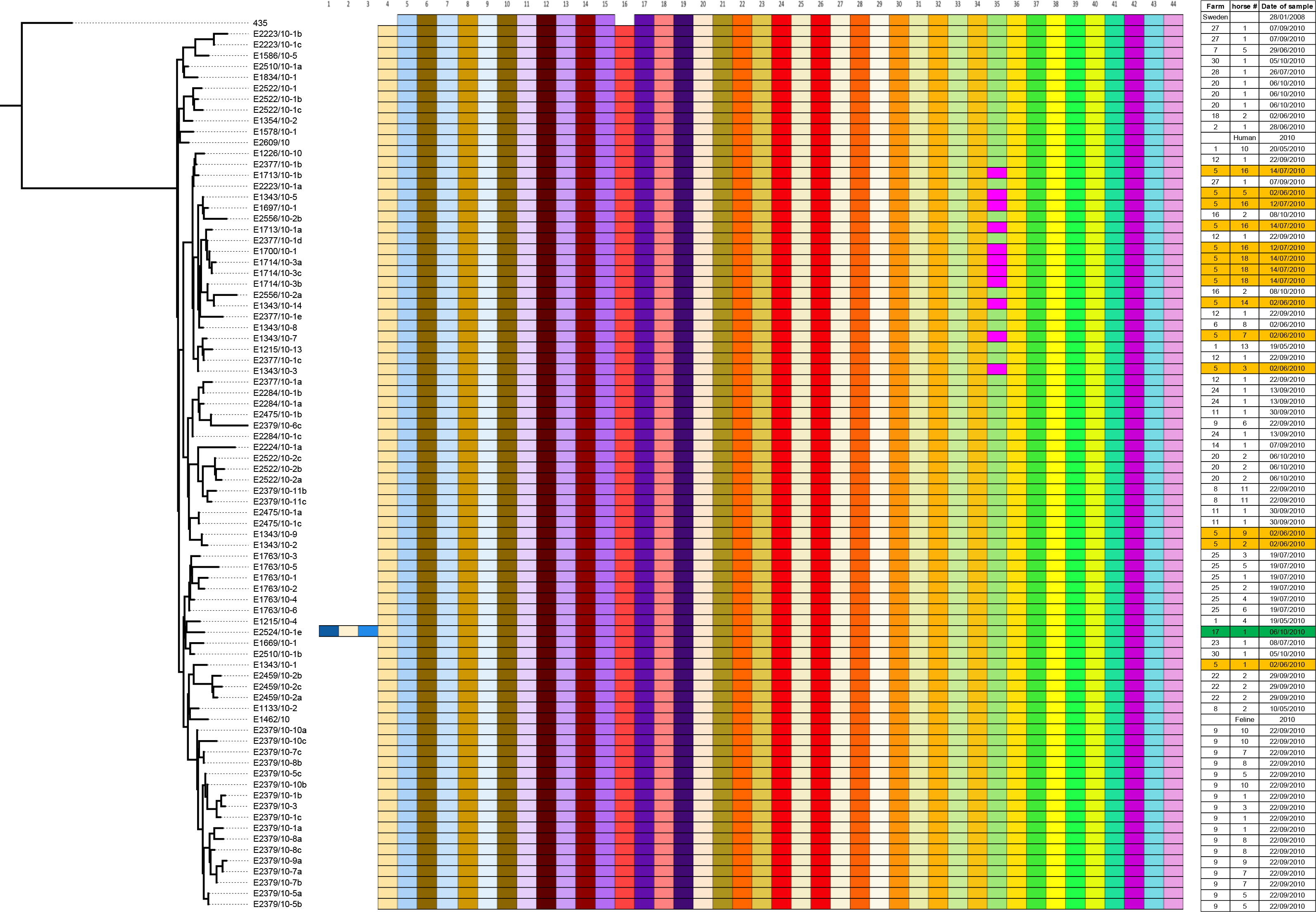
Representation of shared spacer sequences within the CRISPR region of epidemic ST209 isolates relative to the 435 strain that was recovered from Sweden in 2008. The left panel represents the ML phylogeny of *S. zooepidemicus*. The presence of shared spacer sequences, from 1 to 44 is indicated by colored boxes, with the date of isolation, farm and horse number shown in columns to the right. The location of isolates recovered from farm 5 is highlighted by the orange colored entries in the right panel. The isolate from farm 17 that contained an additional three spacers in its genome sequence is highlighted in green in the right-hand panel.

### Epidemiological network analysis identifies an infection source

Network analysis of affected farms identified a single common training yard (yard A) as a primary center of transmission (Figure 3). Two horses leaving yard A on the 19^th^ of February transmitted the disease both to farm 16 and to a stable in the Reykjavik area. All horses (n=20) that left yard A in March, transmitted the disease to new premises (n=18). These new premises each had up to 50 horses stabled and became secondary centers of transmission. However, two horses from yard A, which had returned to farm 16 on the 4^th^ of February, were not incubating the disease. Therefore, it is likely that the transmission of ST209 strains to horses at yard A began between the 5^th^ and 19^th^ of February 2010.

**Figure 3.**
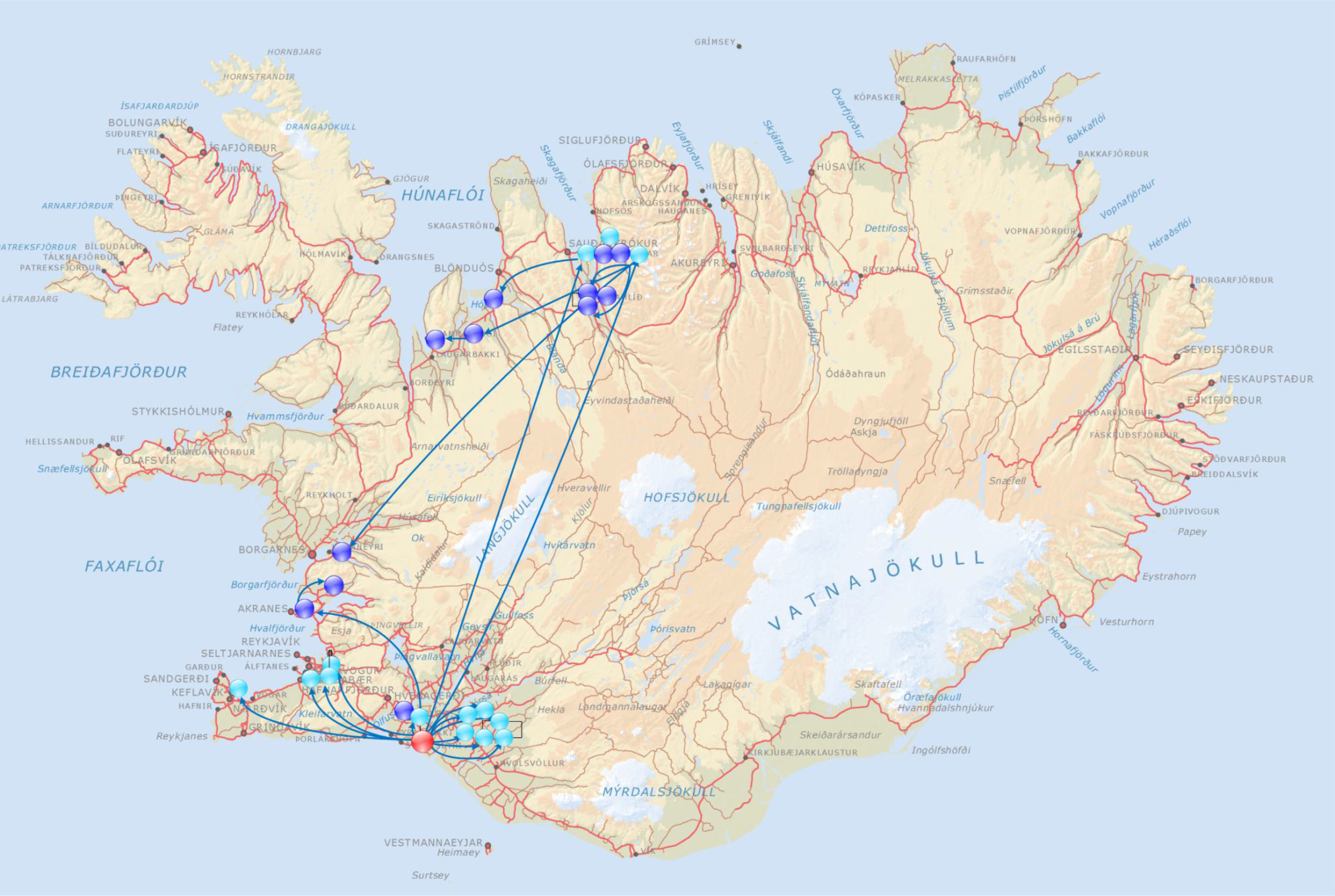
Depiction of the movement of infected horses from the primary center of transmission (red) and the secondary centers of transmission (turkey) to affected farms (blue) during February and March of 2010.

Yard A opened in January 2010 and offers rehabilitation for up to 35 horses recovering from injury or poor condition as well as equipment training for competition horses. Although the epidemic agent was generally transmitted through the coughing of infected horses, coughing was not observed at yard A before April 2010, most probably due to the relatively long incubation period permitting apparently healthy horses to return to their original premises post-training. Instead, the most likely source of transmission of the epidemic strain at yard A was a water treadmill, which horses used on a daily basis. The water used in the treadmill contained no disinfectant and was changed on a once-or twice-weekly basis, providing ideal conditions for the transmission of *S. zooepidemicus* between the visiting horses. Transmission via the water treadmill at yard A provides one possible explanation for the efficiency of the spread of the epidemic throughout Iceland, with multiple secondary sources of infection combining with the original point source to produce a perfect storm of transmission.

### Estimation of the date of common ancestry for the Icelandic ST209 isolates

Following the resolution of the epidemic we hypothesized that the ST209 strain had become endemic within the Icelandic horse population, and that the genetic variation that had accrued since the epidemic could be exploited to estimate the time of most recent common ancestor (TMRCA) shared by the Icelandic ST209 population. To determine if the ST209 strain persisted within the Icelandic horse population, we sampled an additional 36 healthy horses three years after the epidemic. One hundred and seventy-five *S. zooepidemicus* isolates (Supplementary Table 1) were recovered, and of these 20 isolates were identified as ST209, obtained from four healthy horses stabled at different farms throughout Iceland. WGS of the isolates from each of the individual horses indicated that they were very closely related, however, isolates from the different horses clustered separately, with longer branch lengths relative to the isolates recovered from the epidemic in 2010. We analyzed a subset of our dataset, for which the date of isolation was known, with the Bayesian phylogenetics software, BEAST (Drummond and Rambaut 2007). Removal of strains with identical sequences that were recovered from the same animal on the same date produced a dataset of 59 ST209 isolates with 434 polymorphic core genome positions. This dataset was comprised of 4 isolates from 2013, 48 isolates that were recovered during the epidemic in 2010 and 7 non-Icelandic isolates from the collection at the Animal Health Trust, which share an identical or closely related ST to ST209. BEAST includes a number of relaxed molecular clock models that permit modeling of variation in substitution rates on different branches of the tree, allowing correction for the observed rate variation in our data. Utilizing these models, we found that a strict clock with a skyline population model was the best fit of our data and the mean substitution rate per core genome site per year was calculated as 2 × 10^−6^. This substitution rate is similar to core genome rates reported for other streptococci, including *Streptococcus pyogenes* (1.1 × 10^−6^) (Davies, Holden, Coupland, Chen, Venturini, Barnett, Zakour, Tse, Dougan, Yuen et al. 2014) and *Streptococcus pneumoniae* (1.57 × 10^−6^) (Croucher, Harris, Fraser, Quail, Burton, van der Linden, McGee, svon Gottberg, Song, Ko et al. 2011), and many other Gram-positive bacteria, including *Staphylococcus aureus* (3.3 × 10^−6^) (Harris, Feil, Holden, Quail, Nickerson, Chantratita, Gardete, Tavares, Day, Lindsay et al. 2010a). Interestingly, this rate is faster than the closely-related host-restricted pathogen *Streptococcus equi* (5.22 × 10^−7^) (Harris, Robinson, Steward, Webb, Paillot, Parkhill, Holden and Waller 2015) with which *S. zooepidemicus* shares >97% DNA identity. This difference most likely reflects the unusual lifestyle of *S. equi*, which can persist within the guttural pouches of recovered horses (Harris, Robinson, Steward, Webb, Paillot, Parkhill, Holden and Waller 2015). The analysis provided a median estimate for the TMRCA of the Icelandic ST209 strains of July 2008 (95% HPD: August 2007 to May 2009). Our data suggest that the ST209 strains could have circulated in a small number of Icelandic horses prior to the start of the epidemic in 2010. In accordance with our network analysis, the movement of an infected animal to yard A, which received horses from throughout Iceland, was critical for the wider transmission of the ST209 strains. We have previously reported that the within-host diversification of *S. equi* enables individual animals to shed multiple variants of the original infecting strain (Harris, Robinson, Steward, Webb, Paillot, Parkhill, Holden and Waller 2015). Therefore, it is also possible that more than one variant of ST209, potentially originating from the same animal, was introduced to Iceland prior to the start of the epidemic. In this latter scenario, the ST209 variants may have circulated for a shorter period of time within the Icelandic horse population prior to February 2010. However, regardless of the date at which the epidemic strain was introduced to Iceland, the transfer of an infected horse to yard A, identified through our network analysis, remains essential to the rapid spread of the ST209 strains that was characteristic of the epidemic.

### Origins of ST209 strains in Iceland

The Icelandic epidemic ST209 strains share greatest genetic similarity to strain 435, which was isolated from a coughing horse in Sweden during 2008. This strain was selected for sequencing because it shared a common ST209 profile (Supplementary Table 1). To date, with the exception of Iceland, ST209 strains have only been isolated from Scandinavia and it is interesting to note the close relationships between many training yards in Iceland and this region of Europe. The import of horses to Iceland has been prohibited since 1882 for biosecurity reasons. Although the import of used riding equipment is also prohibited, it is difficult to control. Therefore, contaminated tack represents a possible horse-related route of introduction of the ST209 strain.

During the epidemic a ST209 strain was recovered from an infected cat and a lady with septicemia who had suffered a miscarriage that may have been linked to her infection (Figure 4). Previously, a *S. zooepidemicus* strain of ST209 was associated with a zoonotic infection in Finland (Pelkonen, Lindahl, Suomala, Karhukorpi, Vuorinen, Koivula, Vaisanen, Pentikainen, Autio and Tuuminen 2013) and other strains are known to infect companion animals (Webb, Jolley, Mitchell, Robinson, Newton, Maiden and Waller 2008). The ability of ST209 strains to cross host boundaries provides an alternative import mechanism whereby the infection of a human may have facilitated onward anthroponotic transmission to horses resident in Iceland.

**Figure 4.**
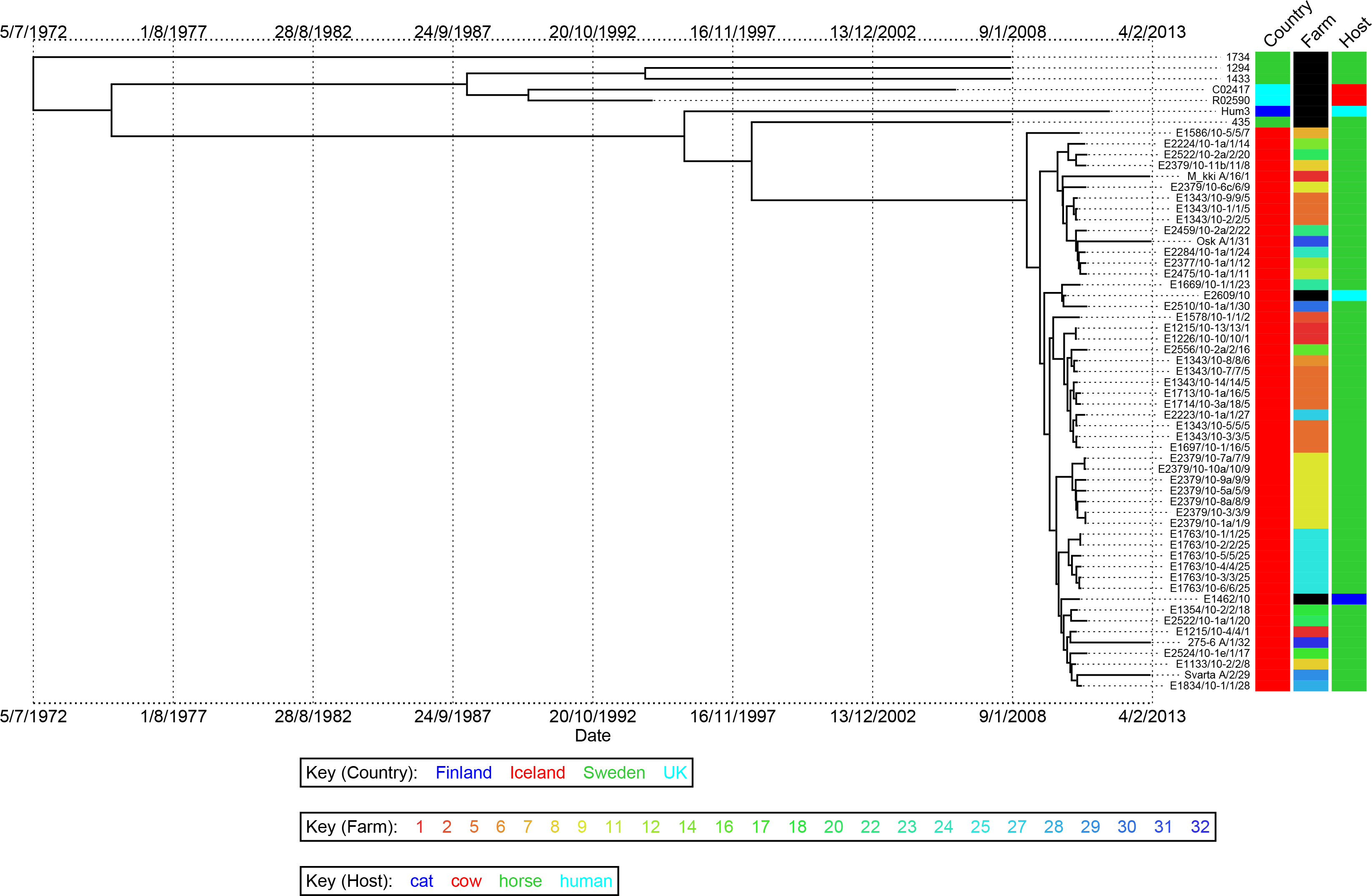
Bayesian phylogenetic visualization of isolate metadata produced with BEAST. Branch lengths represent time with dates shown beneath the tree. Country, farm and host of origin are indicated in columns adjacent to the tree.

## DISCUSSION

By analyzing WGS data of *S. zooepidemicus* isolates collected during the 2010 epidemic of equine respiratory disease, we obtained sufficient data to resolve recent from distant transmission events, and so identify the causal strain. Our data fully support the epidemiological analysis of this epidemic and points to the incursion of a novel strain of *S. zooepidemicus* into the Icelandic horse population. It also reveals the diverse pathogenomic properties of *S. zooepidemicus* and how a novel strain can spread rapidly through a susceptible population devoid of sufficient cross-protective immunity despite a background of concomitant colonization with endemic strains.

This study emphasizes the importance of national biosecurity as a barrier to protect vulnerable populations such as Iceland’s unique horse population.

## METHODS

### Study collection

The origins and details of the isolates of *S. zooepidemicus* that were sequenced in this study are listed in Supplementary Table 1. β-hemolytic colonies of *S. zooepidemicus* strains were recovered from glycerol stocks following overnight growth on COBA strep select plates (bioMérieux). Their identity was confirmed by fermentation of ribose and sorbitol, but not trehalose in Purple broth (Becton Dickinson). The published genome sequences for *S. zooepidemicus* strains MGCS10565 (Beres, Sesso, Pinto, Hoe, Porcella, Deleo and Musser 2008), H70 (Holden, Heather, Paillot, Steward, Webb, Ainslie, Jourdan, Bason, Holroyd, Mungall et al. 2009), BHS5 (Paillot, Darby, Robinson, Wright, Steward, Anderson, Webb, Holden, Efstratiou, Broughton et al. 2010) and ATCC35246 (Ma, Geng, Zhang, Yu, Yi, Lei, Lu, Fan and Hu 2011) were included in the analysis to capture the known species diversity.

### Epidemiological investigation

A questionnaire was sent to 200 premises, including all professional training yards and breeding farms in the country in June 2010 with a follow up in September 2010 to determine whether and when their horses had shown signs of respiratory disease within the preceding months. Further information pertaining to the movement of horses into affected premises was collected by interviews, enabling the network of connected farms and training centers to be investigated.

### Contact study

Three clinically healthy Icelandic horses (horses 16, 17, 18) aged 18-21 years from the Institute for Experimental Pathology at Keldur were transferred to farm 5 two weeks after the first signs of respiratory disease were identified in the resident horses. The experiment was conducted in accordance with the Icelandic animal care guidelines for experimental animals (The Icelandic Animal welfare Act no. 15/1994), the Icelandic Regulations on Animal Experimentation no. 279/2002, which is based on and complies with the European Convention for the Protection of Vertebrate Animals used for Experimental and other Scientific Purposes, European Treaty Series no. 123, 18.III.1986, and after formal approval from the Icelandic ethical committee on animal research, license number 0710-0401. The horses were monitored for the onset of clinical signs by measuring rectal temperature, auscultation of lung and trachea, palpation of submandibular lymph nodes and the pharyngeal area, and assessment of nasal discharge and coughing. Nasal swabs (Coban^®^) for bacteriological and virological examination and blood samples, serum and EDTA (Vacuette) were collected every two days. Horse 17 was euthanized and a post-mortem examination was conducted two days after the first signs of respiratory disease (day 23 post-introduction), where muco-purulent nasal discharge and coughing coexisted with positive cultivation of *S. zooepidemicus*. Horses 16 and 18 were euthanized seven days later and a post-mortem examination was conducted.

### Virus isolation and diagnostics

Nasal swabs taken from horses with clinical signs of respiratory disease were submitted for testing at the Institute of Experimental Pathology in Keldur in Iceland, the National Veterinary Institute in Uppsala, Sweden and the Institute of Virology, Justus-Liebig-Univeritat in Giessen, Germany. Cell free transport media from nasal swabs and enriched peripheral blood leukocyte cells (PBLC)-plasma from EDTA stabilized blood samples collected from the three contact study horses (horses 16, 17 and 18) post-introduction to farm 5 were used to inoculate retroviral vector LXSN116E6E7-transfected foetal lung and kidney cell lines (Thorsteinsdóttir *et al.*, unpublished data). Further samples from swabs taken from several locations in the respiratory tract and conjunctiva post-mortem were also used to inoculate these cells. Inoculated cells were passaged and the conditioned cell culture media was also used to inoculate new cell cultures. The sample was judged as being virus negative if no cytopathic effect could be seen after the third passage. At each passage cells were processed by cytospin and examined by indirect fluorescent antibody staining using pooled convalescent serum samples from 6 horses as the primary antibody. Additional virus isolation attempts were made in the same way with nasal swabs from 14 horses with clinical signs of disease using primary equine foetal lung and kidney cells (Torfason, Thorsteinsdottir, Torsteinsdottir and Svansson 2008), RK-13 cells (ATCC), MA-104 cells (Microbiological Associates) and Vero cells (ATCC).

A wide range of serological tests and PCR assays for equine herpesvirus types 1, 2, 4 and 5, equine infectious anemia virus, equine arteritis virus, equine rhinitis virus A and B, equine influenza A type 1 (H7N7) and type 2 (H3N8), equine salemvirus, human and equine adenovirus, mammalian reovirus serotypes 1, 2 and 3, carnivore parvovirus, human rhinovirus, human enterovirus, canine pneumovirus, influenza A and B, parainfluenza virus 3 and mammalian coronaviruses were used according to the Manual of Diagnostic Tests and Vaccines for Terrestrial Animals produced by the OIE and other published methods (Allard, Albinsson and Wadell 2001; Back, Penell, Pringle, Isaksson, Roneus, Treiberg Berndtsson and Stahl 2015; Balasuriya, Leutenegger, Topol, McCollum, Timoney and MacLachlan 2002; Coggins, Norcross and Nusbaum 1972; Crabb, MacPherson, Reubel, Browning, Studdert and Drummer 1995; Dynon, Varrasso, Ficorilli, Holloway, Reubel, Li, Hartley, Studdert and Drummer 2001; Escutenaire, Mohamed, Isaksson, Thoren, Klingeborn, Belak, Berg and Blomberg 2007; Leary, Erker, Chalmers, Cruz, Wetzel, Desai, Mushahwar and Dermody 2002; Peacey, Hall, Bocacao and Huang 2009; Puig, Jofre, Lucena, Allard, 2Wadell and Girones 1994; Renshaw, Glaser, Van Campen, Weiland and Dubovi 2000; Renshaw, Zylich, Laverack, Glaser and Dubovi 2010; Schunck, Kraft and Truyen 1995; Selvaraju and Selvarangan 2010; Svansson, Roelse, Olafsdottir, Thorsteinsdottir, Torfason and Torsteinsdottir 2009; Thorsteinsdottir, Torfason, Torsteinsdottir and Svansson 2010; Torfason, Thorsteinsdottir, Torsteinsdottir and Svansson 2008) to identify a potential viral cause (Supplementary Table 2).

### Post mortem examinations

A full post mortem examination was performed on the three experimentally infected horses. Samples were taken from all the major organs in addition to samples from the nasal mucosa, larynx and trachea. Tissues were fixed in 10 % neutral buffered formalin. The formalin-fixed material was processed by routine paraffin embedding, and 4 μm thick sections were cut, mounted, and stained with haematoxylin and eosin.

### DNA preparation

A single colony of each *S. zooepidemicus* strain was grown overnight in 3 ml of Todd Hewitt (TH) broth containing 30 μg/ml hyaluronidase (Sigma) at 37 °C in a 5 % CO_2_ enriched atmosphere, centrifuged, the pellet re-suspended in 200 μl Gram +ve lysis solution (GenElute, Sigma) containing 250 units/ml mutanolysin and 2 × 10^6^ units/ml lysozyme and incubated for 1 hour at 37 ^o^C to allow efficient cell lysis. DNA was then purified using GenElute spin columns according to manufacturer’s instructions (all Sigma).

### MLST assignment

MLST sequence types were identified from sequence data as previously described (Webb, Jolley, Mitchell, Robinson, Newton, Maiden and Waller 2008).

### DNA Sequencing

DNA libraries for isolates recovered pre-2011 were created using a method adapted from the Illumina Indexing standard protocol. In brief, the steps taken (with clean-up between each step) were: genomic DNA was fragmented by acoustic shearing to enrich for 200 bp fragments using a Covaris E210, end-repaired and A-tailed. Adapter ligation was followed by overlap extension PCR using the Illumina 3 primer set to introduce specific tag sequences between the sequencing and flow-cell binding sites of the Illumina adapter. After quantitation by qPCR followed by normalization and pooling, pooled libraries were sequenced on Illumina GAII and HiSeq platforms according to the manufacturer’s protocols generating index tag-end sequences. For isolates recovered post-2011, DNA libraries were prepared using the Illumina Nextera XT DNA library prep kit, according to the standard protocol. The DNA libraries were pooled and quantified using the KAPA Library Quantification Kit for Illumina platforms prior to sequencing on Illumina MiSeq according to the manufacturer’s protocol.

### Variation detection

Illumina reads were mapped onto the relevant reference sequences using SMALT (http://www.sanger.ac.uk/resources/software/smalt/); H70 (accession number FM204884) for the total population, and a de novo assembly of Clade 1. A minimum of 30x depth of coverage for more than 92 % of the reference genomes was achieved for both references (Supplementary Table 1). The default mapping parameters recommended for reads were employed, but with the minimum score required for mapping increased to 30 to make the mapping more conservative. Candidate SNPs were identified using SAMtools mpileup (Li, Handsaker, Wysoker, Fennell, Ruan, Homer, Marth, Abecasis and Durbin 2009), with SNPs filtered to remove those at sites with a mapping depth less than 5 reads and a SNP score below 60. SNPs at sites with heterogeneous mappings were filtered out if the SNP was present in less than 75 % of reads at that site (Harris, Feil, Holden, Quail, Nickerson, Chantratita, Gardete, Tavares, Day, Lindsay et al. 2010b). Identification of the core genomes was performed as previously described (Harris, Feil, Holden, Quail, Nickerson, Chantratita, Gardete, Tavares, Day, Lindsay et al. 2010b; Holden, Hsu, Kurt, Weinert, Mather, Harris, Strommenger, Layer, Witte, de Lencastre et al. 2013). Recombination was detected in the genomes using Gubbins (http://sanger-pathogens.github.io/gubbins/) (Croucher, Page, Connor, Delaney, Keane, Bentley, Parkhill and Harris 2014). Phylogenetic trees for ST209 and ST22 were constructed separately using RAxML v7.0.4 (Stamatakis 2006) for all sites in the core genomes containing SNPs, using a GTR model with a gamma correction for among site rate variation (Harris, Feil, Holden, Quail, Nickerson, Chantratita, Gardete, Tavares, Day, Lindsay et al. 2010b). We used the Bayesian software package BEAST (v1.7.4) (Drummond, Suchard, Xie and Rambaut 2012) to investigate the temporal, spatial and demographic evolution of the clade 1 population. To estimate the substitution rates and times for divergences of internal nodes on the tree, a GTR model with a gamma correction for among-site rate variation was used. All combinations of strict, relaxed lognormal, relaxed exponential and random clock models and constant, exponential, expansion, logistic and skyline population models were evaluated. For each, three independent chains were run for 100 million generations, sampling every 10 generations. On completion each model was checked for convergence, both by checking ESS values were greater than 200 for key parameters, and by checking independent runs had converged on similar results. Models including logistic population models failed to converge so were discarded. Models were compared for their fit to the data using Bayes Factors based on the harmonic mean estimator as calculated by the program Tracer v1.4 from the BEAST package. The best-fit model was found to be a strict skyline population model. A maximum clade credibility (MCC) tree was created from the resulting combined trees using the treeAnnotator program, also from the BEAST package.

### ST209-specific qPCR

ST209 isolates (clade 1) were identified using a specific qPCR for the *nrdE* allele 31. Forward (5’-ACCAAAAAAGAAAATGCT-3’) and reverse (5’-TCAACACTATAAGGACTAAAGAGA-3’) primers were used to amplify the *nrdE* allele 31 on a STEPONE plus machine (Applied Biosystems) with KAPA SYBR FAST ABI PRISM reagents (Kapa Biosystems) and cycling conditions of 95 °C for 3 minutes followed by 40 cycles of 95 °C for 30 seconds and 62 °C for 10 seconds. A melt curve was performed between 60 °C and 95 °C reading SYBR every 0.3 °C. All qPCR experiments were performed in triplicate.

## DATA ACCESS

The Illumina sequences generated and used in this study have been deposited in the European Nucleotide Archive under the study accession number ERP000883. The accession numbers for the sequences of each isolate are listed in Supplementary Table 1.

## ACKNOWLEDGEMENTS

The authors thank Dr. Louise Treiberg Berndtsson and co-workers at the National Veterinary Institute in Uppsala, Sweden and Dr. Christine Förster at the Institute of Virology, Justus-Liebig-Univeritat in Giessen, Germany for their assistance with virological testing and the personnel at the department of bacteriology at Keldur for their technical assistance. The Icelandic human isolates were obtained from the Department of Clinical Microbiology, Landspítali University Hospital, Reykjavik, Iceland. The human isolates from the United Kingdom were provided by Professor Androulla Efstratiou at Public Health England. Bovine isolates were provided by Dr. Adrian Whatmore of the Animal and Plant Health Agency. The human isolate from Finland was a kind gift from Professor Sinikka Pelkonen and Dr. Tamara Tuuminen of the Finnish Food Safety Authority and University of Helsinki, respectively. We are grateful to Dr. Alistair Darby of the University of Liverpool for providing the genome sequence of *S. zooepidemicus* strain 1175. We also thank the core sequencing and informatics teams at the Sanger Institute for their assistance and The Wellcome Trust for its support of the Sanger Institute Pathogen Genomics and Biology groups. SRH, JP and MTGH were supported by Wellcome Trust grant 098051.

## AUTHOR CONTRIBUTIONS

SB, VS, EG, OGS, JRN, MTGH and ASW designed the study. SB, SRH, VS, EG, OGS, KG, KFS, CR, ARLC, MTGH and ASW carried out the research. SB, VS, EG, OGS, JP and ASW supplied isolates, metadata and whole genome sequencing. SB, SRH, VS, EG, KFS, JRN, CR, ARLC, MTGH and ASW analyzed the data. SB, SRH, VS, EG, OGS, MTGH and ASW wrote the manuscript. All authors read and approved the final manuscript.

